# Genetic Variation in Honey Bee Queen Reproductive Performance: Implications for Colony Growth and Sustainability

**DOI:** 10.1101/2025.05.09.653176

**Authors:** Zhaorigetu Hubhachen, Heaven Strachan, Xaryn Cleare, Danielle Kroh, Hongmei Li-Byarlay

## Abstract

The reproductive performance of honey bee (*Apis mellifera* L.) queens is a key determinant of colony health and productivity. However, variation in queen fecundity among different genetic lineages remains poorly understood. In this study, we compared the egg-laying capacity and colony demographics of queens from three honey bee populations, Russian, feral, and commercially reared stocks over multiple years (2020, 2021, and 2024). Using standardized colony conditions and controller assessments of queen egg-laying rates, we found that Russian and feral queens exhibited significantly higher fecundity than commercial queens. In 2024, we further observed that worker populations in Russian and feral colonies increased over time whereas commercial colonies experienced a decline, suggesting potential differences in colony sustainability. While no significant differences were detected in pollen storage or brood area in earlier years, trends in brood production alighted with egg-laying patterns. Our results highlight the importance of genetic background in shaping queen reproductive performance and colony growth. These findings provide insights for breeding programs aimed at improving queen quality and colony resilience, particularly in the face of environmental stressors such as *Varroa* infestation and resource availability.

## Introduction

Honey bee (*Apis mellifera* L.) colonies function as complex superorganisms, where division of labor and social cohesion are essential for survival (Winston 1987; Amdam and Page 2010). Within this social structure, the queen bee is holding the reproductive capacity and in charge of colony growth, worker productivity, and overall resilience (Walsh et al. 2020). Queen fecundity determines the availability of workers for foraging, brood care, and defense, thereby influencing critical colony functions (Kocher et al. 2009). However, not all queens exhibit the same reproductive performance, and variation in egg-laying rates may have profound consequences for colony stability. Understanding how genetic background influences queen fecundity is essential for evaluating colony fitness and long-term viability.

Social dynamics within a colony are heavily influenced by queen-worker interactions (Winston 1987). Worker bees regulate brood-rearing activity based on the queen’s reproductive output, responding to shifts in egg-laying rates by adjusting foraging effort, food storage, and nursing behavior (Walsh et al. 2020). Colonies experiencing reproductive instability, whether due to poor queen quality or external stressors, may exhibit behavioral changes such as supersedure, decreased brood care, or increased aggression (Amiri et al. 2017). These responses can impact colony cohesion and may even contribute to colony collapse if worker populations decline too rapidly (Tarpy et al. 2023). The interplay between queen reproductive success and worker behavioral plasticity highlights the importance of selecting high-quality queens for sustainable beekeeping and conservation.

Ecological stressors, including habitat loss, pesticide exposure, and pathogen pressure, exacerbate the challenges faced by honey bee colonies (Mullin et al. 2010; Traynor et al. 2016; Wu-Smart and Spivak 2016; Kulhanek et al. 2017; Ma et al. 2019; Wen et al. 2021; Wang et al. 2022; Siviter et al. 2023). One of the most significant threats is *Varroa destructor*, an ectoparasitic mite that weakens colonies by feeding on developing brood and vectoring viral infections (Le Conte et al. 2010; Guichard et al. 2023; Zheng et al. 2023). Certain honey bee populations, such as Russian and *Varroa* Sensitive Hygiene bees, have shown greater resistance to *Varroa* infestations and associated pathogens, raising the question of whether these populations also exhibit superior reproductive performance (Ibrahim et al. 2007; Rinderer et al. 2010; Unger and Guzman-Novoa 2010; Danka et al. 2011; Tsuruda et al. 2012; Kirrane et al. 2015; Guichard et al. 2023). Feral colonies also displayed resilience to parasite and mite resistance from previous reports around the world and in North America (Seeley 2007; Hunt et al. 2016; López-Uribe et al. 2017; Oddie et al. 2017; Belsky and Joshi 2019; Morfin et al. 2020; Russo et al. 2020; Hinshaw et al. 2021; Smith et al. 2021; Ward et al. 2022; Guzman-Novoa et al. 2024). If resistant populations maintain a higher queen fecundity, they may be better equipped to sustain worker numbers and recover from environmental stressors. This study explores whether genetic variation among Russian, feral, and commercially reared queens influences colony reproductive success and social stability.

The quality of a queen bee, her reproductive capacity, and overall colony demographics are key indicators of colony health (Lee et al. 2019; Tarpy et al. 2023). Several methods exist for assessing queen quality, with one of the most direct measurement of egg-laying rate is to record the number of eggs laid by a queen during active seasons (Walsh et al. 2020). Successful fertilization of worker-destined eggs depends on the viability of sperm stored in the queen’s spermatheca, a crucial factor influencing reproductive success (Baer et al. 2016). In addition to egg-laying rates, queen quality can be evaluated through traits such as brood pattern consistency, body weight and morphology, ovary development, sperm viability, temperament, longevity, and disease resistance (Amiri et al. 2017; Scaramella et al. 2023). However, due to the scope of this study and constraints imposed by the COVID-19 pandemic, we focused exclusively on egg-laying performance and colony demographics in brood area, pollen storage and number of worker bees. They serve as fundamental indicators of queen quality and may provide insights into additional reproductive and colony health parameters in future studies.

To investigate these relationships, we compared egg-laying rates and colony demographics across three genetic stocks: Russian, feral, and commercial colonies. We examined whether variations in queen fecundity correlated with worker population size, brood area, and pollen storage over multiple years. By quantifying these colony-level traits, we aimed to determine whether differences in queen reproductive capacity translate into long-term colony resilience. Given the importance of worker population dynamics in colony survival, understanding these genetic influences could inform breeding programs focused on improving honey bee health and sustainability.

## Materials and Methods

### Colonies and queens

Queens used for the study were from three lines: Russian stocks, feral bees, and commercially reared bees. The Russian honey bee queens used in the experiments were same year grafted queens from a single source provided by the United States Department of Agriculture, Agricultural Research Service (USDA ARS) in Baton Rouge, Louisiana. The feral honey bee queens were grafted from feral colonies in West and Central Ohio. These colonies were captured by swarm traps in remote areas, specifically in tree cavities in natural habitat. Based on reports from beekeepers and our own observations, all feral colonies had survived naturally for at least two winters before we collected them for this study. As for the commercially reared colonies, queens were grafted from package or commercially available colonies purchased from Combs Bee Farm in Urbana, Ohio. The farm had imported the bees from a commercial bee farm in Georgia, USA. In both 2020 and 2021, we set up three colonies/genetic background/year (*n*=3) whereas in 2024, we set up six colonies/genetic background (*n*=6). All queens underwent grafting and shared identical environmental conditions during their development. They were permitted to engage in open mating in June of each year. All the grafted queens originated from a same mother colony, each possessing distinct genetic backgrounds (Russian, Feral, or Commercial).

Our experiment commenced with queens of the same age with all being one-year-old. These queens were introduced into 5-frame colonies with frames measuring 9_⅛_ inches (23.18 cm) in depth. Each nucleus included a minimum of two capped brood frames, two frames stocked with food (honey and pollen), and a frame of pre-established empty combs. The bee populations in these nuclei were sourced from existing colonies with similar genetics background. Each nucleus began with an equal workforce and followed identical management protocols. Consistency in foraging environments was maintained for all colonies, with uniform weather and temperature conditions.

All the colonies involved in the study were established within our research apiary in Yellow Springs, Ohio, with GPS coordinates of 39.789825103687065 and -83.92233587174182. We consistently monitored the conditions of both the bees and each queen. Data on colony demographics were collected, including adult bee populations on each frame, the brood area or the percentage of each frame, and the pollen area or percentage of each frame, as illustrated in Fig. 1. To evaluate the egg-laying capabilities of the queens, we employed Jenter kits, as depicted in Fig. 1B. In 2020 and 2021, queens were given a period of 3-4 weeks in their new nuclei to acclimate before being caged.

**Fig. 1.**
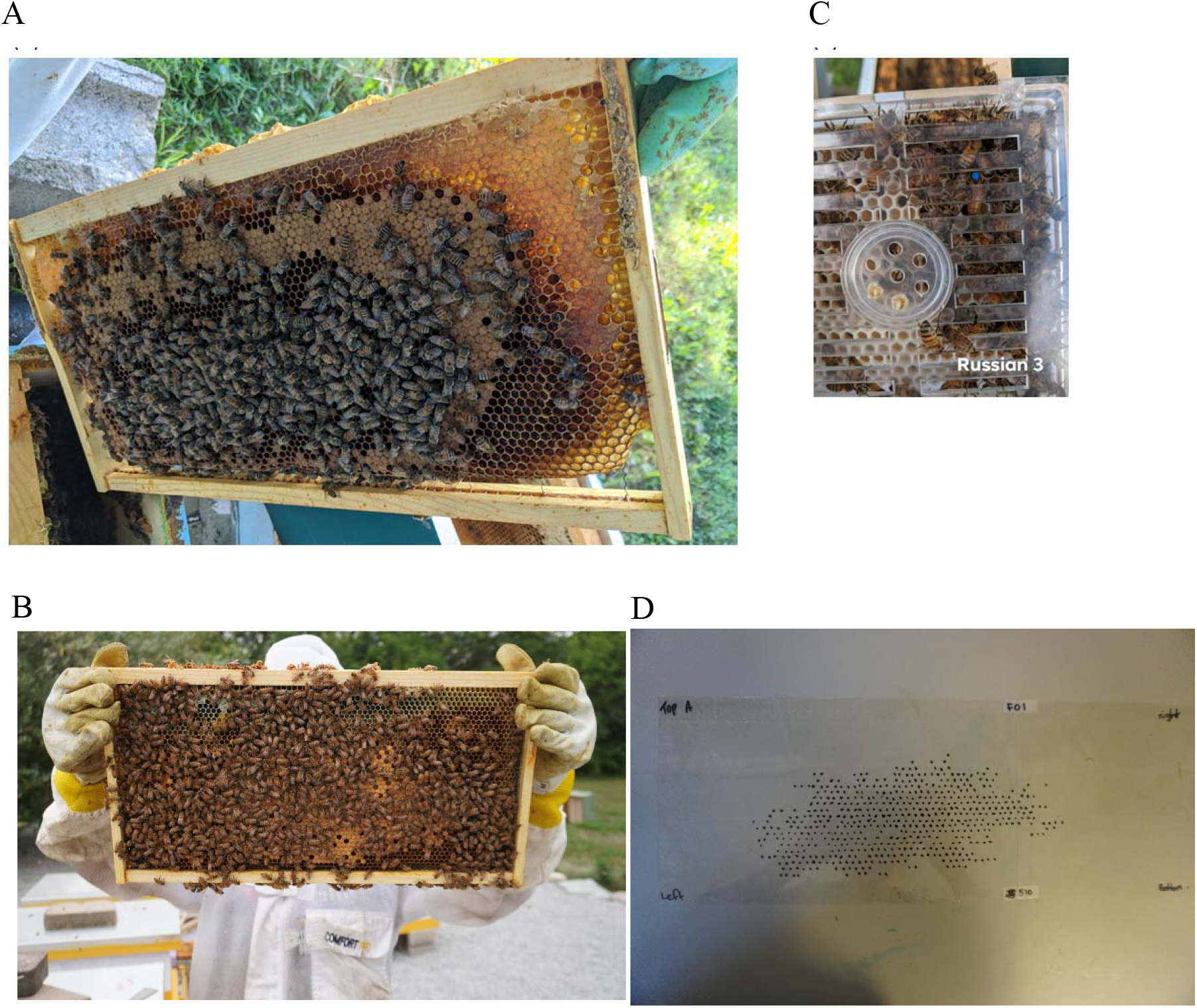
Colony frames and eggs counting. **A:** An example of a colony frame used to record the colony demographics. **B:** An example of a colony frame used to record the worker numbers in 2024. **C:** One Jenter kit was used to cage each queen. The queen was marked with a blue color. **D**: eggs were marked using a transparent plastic sheet in 2024.

All queens were able to lay eggs regularly in the combs before being tested in the Jenker kits (Fig. 1C). All the participating colonies were tested in July 2020 and 2021. We allowed the existing workforce to be replaced by the new worker workforce of the new queens. In the kit, each queen could lay up to 112 eggs. Queens were caged for 24 hours (once a week for 3 weeks), and the number of eggs was recorded from each Jenter kit. These recordings were repeated in the following 2 weeks. We ensured that each queen adapted and laid eggs in the new Jenter kit, and took measures to prevent worker bees from removing the eggs from the cells in the Jenter kit. Queens that failed to lay any eggs were excluded from the experiment.

In 2024, commercial and Russian mated queens were purchased from Combs bee farm, Ohio and Coy bee farm, Mississippi, respectively whereas mated feral queens were obtained from Young Bee Farm, KY. To measure the natural egg-laying capacity of queens, we provided a full frame for egg-laying. The experiment initially included 30 queens: 12 commercial, 8 Russian, and 10 feral. By the conclusion of the study, we successfully collected complete data from 6 queens in each category. Each hive was configured with two to three boxes, depending on colony size. The bottom boxes contained brood, eggs, larvae, and pollen, while a queen excluder was kept between the lower and upper boxes. The top box contained foundation frames with no drawn comb or fully capped honey frames, ensuring queens had no other available space for egg-laying. A single empty frame with drawn comb was kept in the center of the top box and a queen was introduced in the top box of each hive before sealing it. We removed the previously drawn-out frame and placed it in a freezer for later eggs counting. Then a new empty frame was introduced into the top box and allowed the queen to lay eggs. After 24h, the frame was removed and stored as described previously. The queen was then released back into the bottom boxes for a week. After this period, the queen was removed back to the top box, and the newly laid egg frames were collected. This process was repeated to obtain four replicates, with each queen laying eggs within 24h period per replicate. The experiments were conducted between June and July of 2024.

### Eggs frame reading

To count the eggs accurately, we prepared a same size of transparency sheet for each frame and taped onto it. Then we marked each cell with egg by sharpie under a flashlight/head lamp (Fig. 1C). We estimated the numbers of brood and workers with following the method described by Simone-Finstrom et al. (2016). The number of workers, pollen storage areas, and brood proportions were estimated by taking photographs of each colony.

### Data analysis

The demographic data, which included the counts of adult bees, pollen storage areas, and brood areas were subjected to analysis using ANOVA to assess variations among frames from each colony. A one-way analysis of variance (ANOVA) was employed to investigate differences in fecundity means. In the statistical model, colony and location were treated as random effects. Tukey post hoc tests were conducted to perform pairwise comparisons between the various experimental groups. Statistical significance was determined at α ≤ 0.05.

## Results

We compared the colony demographics including brood percentage, pollen storage, and number of adult bees per frame. For brood populations, we measured the percentage of brood area (sealed brood areas) of each queen per observation. No significant difference was found in the percentage of sealed brood area per observation among the three lines (Fig. 2A) (*F*_2, 46_ = 0.32; *p* = 0.72). However, the percentage of brood exhibited higher in both Russian (40.71%) and feral (38.68%) lines than that of commercially reared bees (32.19%).

**Fig. 2.**
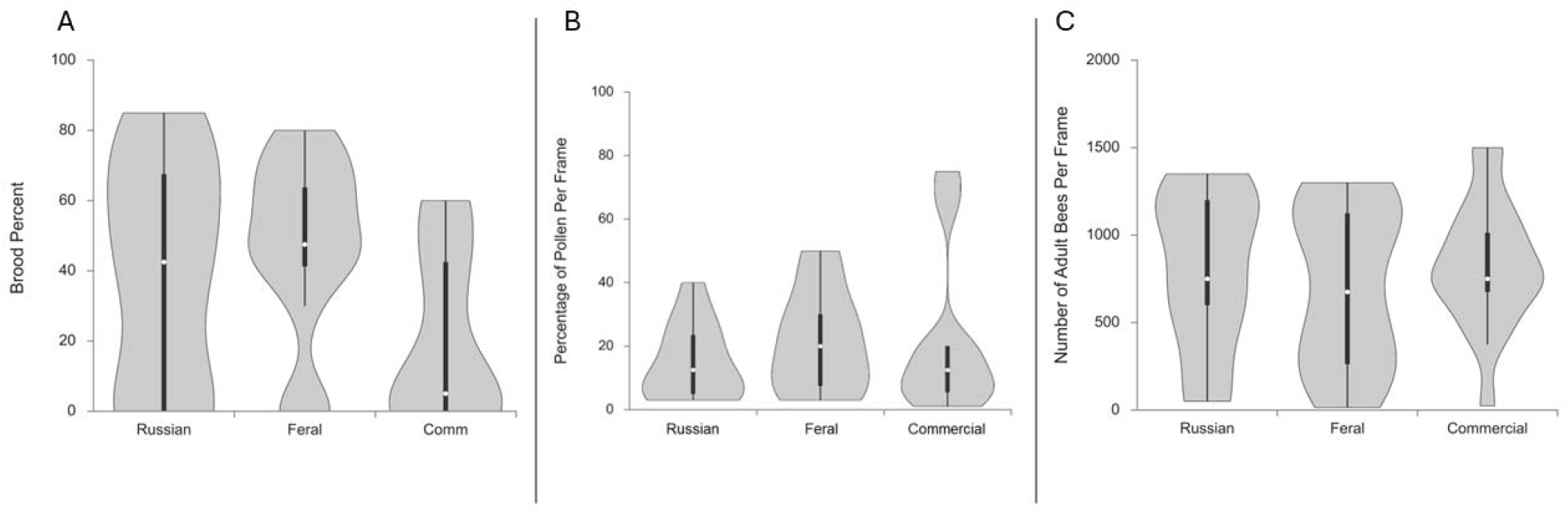
Colony demographics from 2020-2021. **A:** Violin plot displaying the percentage of brood (pupa) per frame in each tested colony. A significant difference was found among the three genetics (ANOVA, *p* = 0.02). A post hoc Tukey test showed that the colonies of feral lines have a higher percentage of sealed brood compared to that of the commercially reared colonies (Tukey-Kramer HSD: Q = 3.99; *p* < 0.05). **B:** Violin plot displaying the percentage of pollen per frame in each tested colony. No significant difference was found among the three treatments (ANOVA, *p* = 0.60). **C**: Violin plot displaying the total number of adult worker bees in each colony tested in 2020-2022. No significant difference was found among the three treatments (*p* = 0.65). The boxplots are shown inside the violin.

To investigate the pollen storage of each line, we compared the area of pollen and bee bread within each colony in 2020 and 2021. No significant difference was found among the colonies for the average percentage of cells containing pollen or bee bread (*F* _2, 31_ = 0.52; *p* = 0.60). The average percentage of cells containing pollen in the Russian, feral, and commercially reared colonies was 14.92%, 15.08%, and 22.30%, respectively (Fig. 2B).

For adult worker populations, we estimated the number of adult workers for each queen per observation. No significant difference was found in the number of adult workers per observation among the three lines in 2020 and 2021 (Fig. 2C) (*F* _2, 53_ = 0.43; *p* = 0.65). The average number of adult workers in the Russian, feral, and commercially reared colonies was 789.71 ± 110.35, 710.00 ± 104.15, and 834.50 ± 80.50, respectively.

To compare queen egg-laying behavior, we measured the mean number of eggs laid by each queen. A significant difference was found in queen egg-laying rates per observation among the three lines in 2020 and 2021 (*F* _*2*,15_ = 41.61; *p* <0.001) (Fig. 3 and Supp. Table S1-S4).

**Fig. 3.**
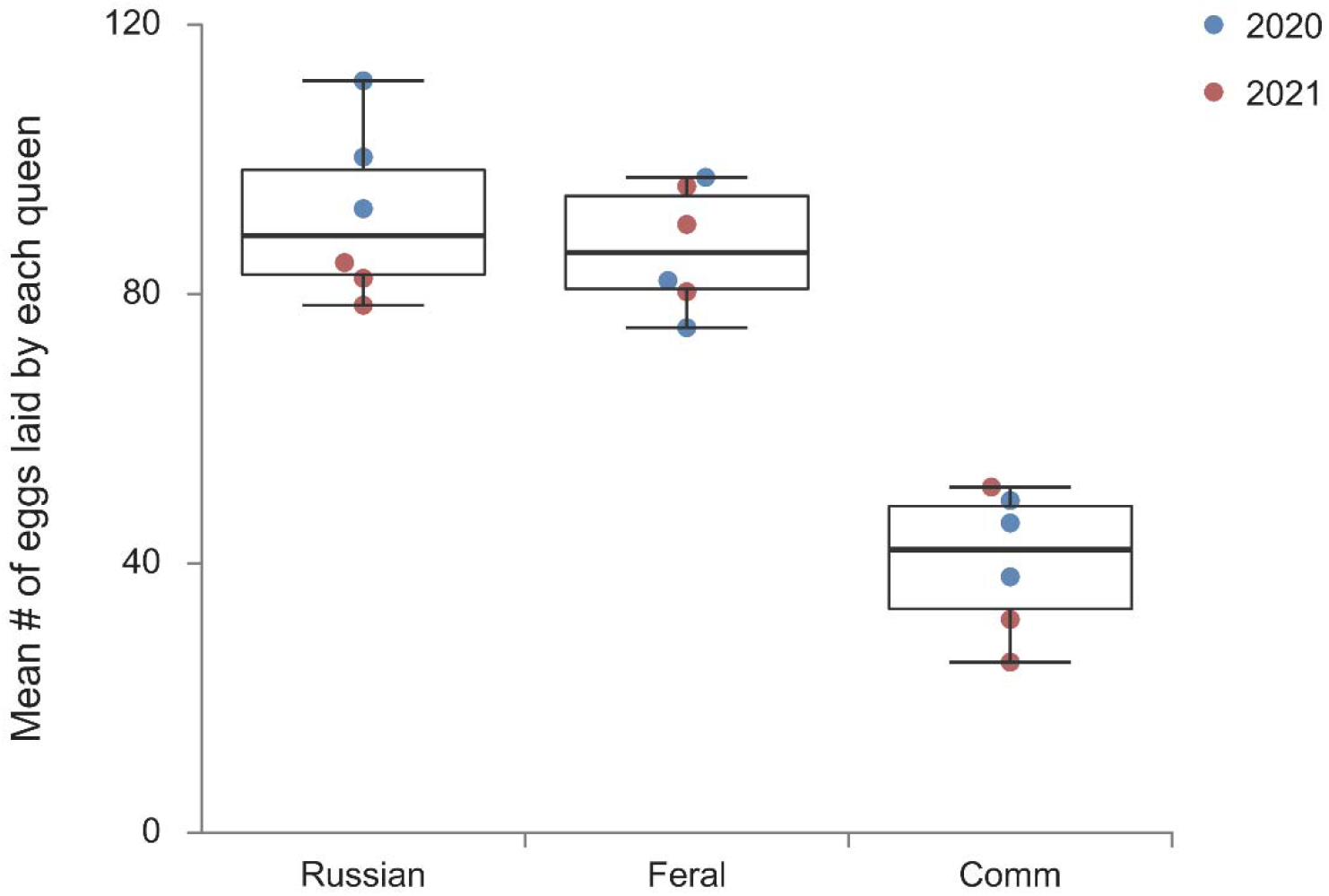
Boxplot displaying the mean number of eggs laid per queen from 2020-2021 experiment. Three genetics of populations (Russian, Feral, and Commercial) were tested. A significant difference was found in queen egg-laying rates per observation among the three lines (*p* < 0.001) (Supp. Table S1 - 4). A post hoc Tukey test showed that queens from the Russian and feral lines laid more eggs compared with queens in the commercially reared colonies (Tukey-Kramer HSD: Q = 11.68; *p* < 0.01). Queens from the feral lines also laid more eggs compared with queens in the commercially reared colonies (Tukey-Kramer HSD: Q = 10.58; *p* < 0.01). There was no significant difference between Russian and feral bees (Tukey-Kramer HSD: Q = 1.10, *p* = 0.71). Each circle was a data point.

In 2024, we estimated the worker numbers at the beginning and the end of the experiment. The estimated worker numbers in the commercial colonies were decreased but not significantly (*p*=0.26) whereas those in the Russian (*p*=0.31) and Feral (*p*=0.03) colonies were increased (Fig. 4A). The estimated worker numbers in the beginning in commercial, Russian and feral colonies were 14,456 ± 1,245.37, 9,778 ± 1,075.23, and 9,278 ± 981.40, respectively whereas those in the end in these colonies were 12,284 ± 1142.95, 11,295 ± 727.07, and 13,019 ± 941.82, respectively.

**Fig. 4.**
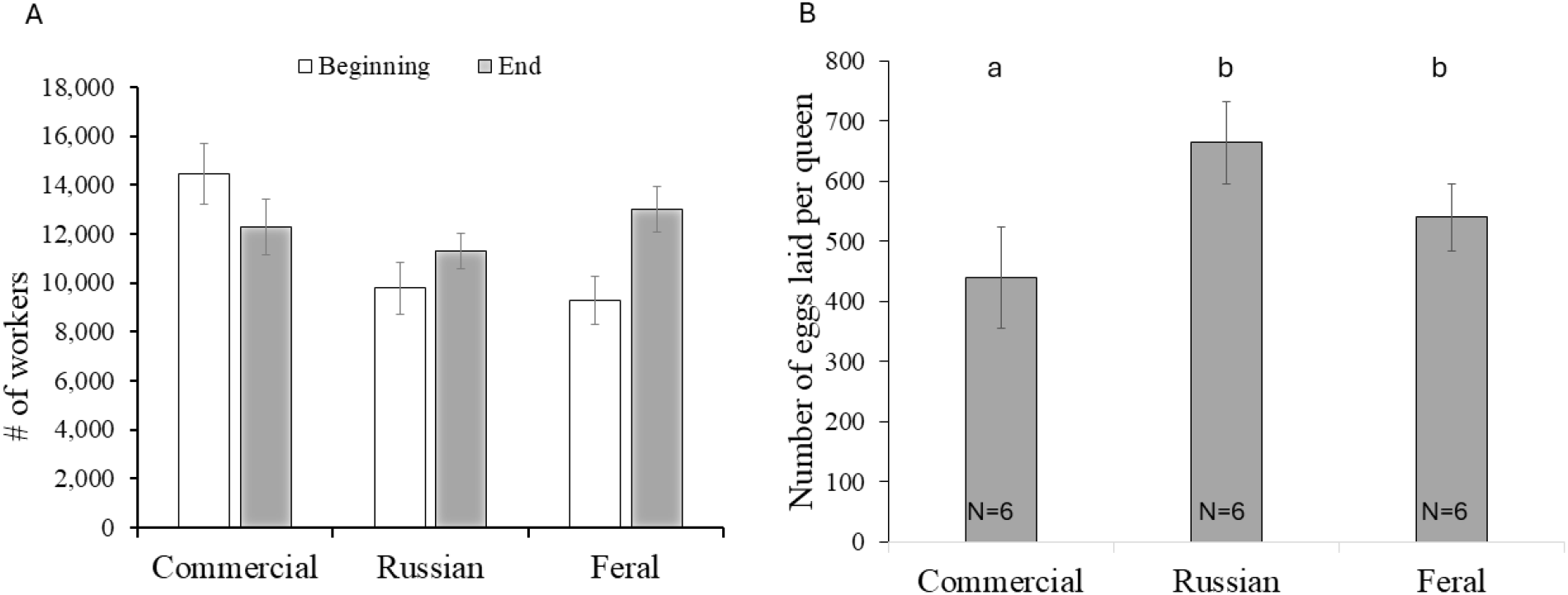
Bar graph showing (A) the total number of workers at the beginning and end of the experiment, (B) the mean number of eggs laid per queen in 2024. The number of queens of each genotype is 6. Total there were 18 queens being tested. Each queen was tested for 4 replicates.

To allow queens to be freely walking in a natural setting, we used the frames to let queens lay eggs in all three colonies in 2024. A post hoc Tukey test showed that queens from both Russian and feral lines laid significantly more eggs than the queens in the commercial colonies (Fig. 4B, Q = 11.68; *p* < 0.01, and Q = 10.58; *p* < 0.01). There was no significant difference between Russian and feral bees (Q = 1.10, *p* = 0.71).

## Discussion

Our results compared colony demographics and queen fecundity across three different lines of honey bees: Russian line, feral bees, and commercially reared bees. The primary outcome revealed that queens from the Russian lines and feral bees exhibited much higher egg-laying capacities compared with queens from commercially reared bees. Based on the colony demographics data, adult bee numbers and pollen storage did not show a significant difference, but the brood pattern shows the same trend as the egg-laying data. This discovery highlights the queen quality of different bee genetics, which provided the potential for bee breeding (Kirrane et al. 2015, Guichard et al. 2023). In addition, our results show that the worker populations were declining trend in commercial colonies, whereas they were increasing in both Russian and feral colonies when estimated them at the beginning and end of the experiment. The population decline in the commercial colonies was most likely due to the infestation of varroa mites as reported by other studies (Woods, 2021; Insolia et al., 2022; Warner et al., 2024). For example, Woods (2021) and Insolia et al. (2022) reported that up to 43% of honeybee colonies lost due to *Varroa* mite infestation from April 2019 to April 2020.

Our study provides novel insights into the ecological and evolutionary dynamics of queens’ egg-laying behavior in honeybee colonies. By integrating behavioral observations with quantitative analyses, we demonstrate that genetics significantly influence queen reproductive strategies, ultimately shaping colony-level fitness outcomes. The findings underscore the adaptive plasticity of queen egg-laying rates in response to colony demography and environmental conditions. Queens modulate their reproductive output in accordance with worker availability and resource abundance. This aligns with life-history theory, which predicts that reproductive effort should be adjusted based on ecological pressures to maximize lifetime fitness.

Moreover, our results suggest that variation in egg-laying rates may serve as a mechanism of reproductive conflict resolution within colonies. Differential reproductive investment by queens in response to worker policing and brood viability signals a sophisticated feedback system that maintains colony homeostasis. Egg-laying behavior is a primary indicator of queen quality in honey bee’s colony health (Amiri et al., 2017). Queens’ oviposition and their resistance to diseases are important to study for colony survival (Amiri et al., 2017; McAfee et al., 2020). Additionally, maintaining genetic diversity is essential for enhancing a colony survival and performance while reducing the risk of inbreeding (Tarpy et al., 2023). Future research should explore the genetic and physiological underpinnings of this dynamic, particularly how hormonal regulation mediates reproductive decision-making in eusocial insects.

Our study highlights the role of worker interactions in shaping queen reproductive behavior. Queens in larger colonies exhibited higher egg-laying rates, suggesting that worker density and brood care capacity provide key regulatory cues. This supports previous findings that worker feedback, including pheromonal signaling and trophallaxis, influences a queen reproductive strategy (Moore et al., 2015).

Brood patterns are also critical to measure the queen quality. A brood pattern is considered poor if more than 20% of the cells of the sealed brood are empty. This indicates that either the queen is not laying enough eggs or the developing bees are not surviving to emerge. Besides assessing queen quality, the colony environment can influence the brood pattern. Poor brood patterns have been associated with pathogenic infections such as microsporidians, bacteria, viruses, or fungi. In addition, some heritable worker traits, such as the removal of diseased or *Varroa*-infested brood, may result in a worse brood pattern but contribute to a healthier colony (Borba et al., 2022). Our data indicates that there is a difference in the brood pattern between the feral colonies and commercial colonies.

The ability of queens to dynamically adjust their egg-laying rates has profound consequences for a colony fitness. Optimal reproductive allocation ensures a balance between workforce maintenance and brood production, crucial for colony survival and competitive success. Our study contributes to a broader understanding of how intrinsic and extrinsic factors interact to shape reproductive strategies in eusocial insects.

Given the ongoing threats to pollinator populations, understanding the mechanisms underlying reproductive plasticity in honey bees is critical. Conservation efforts should consider how genetics and environmental pressures influence queen fecundity and, consequently, colony persistence (Wu-Smart and Spivak 2016).

Our study has contributed novel insights into the significance of egg-laying behavior and brood patterns as pivotal markers of queen quality within the context of three different bee genetic lines. The knowledge gained from the present study holds promise as a valuable tool for identifying and selecting high-quality queens in breeding and selection programs. A healthy and well-mated queen can produce a workforce of genetically diverse and robust worker bees, which enhances the colony’s resilience to environmental challenges and threats such as *Varroa* mites, pathogens, and pesticides.

This study enhances our understanding of honey bee reproductive ecology by demonstrating how queen egg-laying rates are shaped by genetics. We provide a foundation for future studies into adaptive significance of reproductive plasticity in eusocial insects. Continued investigations from molecular and physiological perspectives will be essential for mitigating the challenges of pollinator deficit problems and ensuring the stability of colonies.

## Acknowledgments

We thank Drs. Lilia de Guzman and Michael Simone-Finstrom for their help at the USDA ARS Baton Rouge Laboratory to provide the Russian queens. We also thank Ashley Cordle’s assistance in the field and CSU bee lab members’ input to improve the manuscript. Funding: Financial support was granted to HL-B by the United States Department of Agriculture (USDA), National Institute of Food and Agriculture (NIFA) Evans-Allen fund (Grant Number NI241445XXXXG004), a USDA Sustainable Agriculture Research and Education North-Central Region Partnership Grant (Grant Number ONC19-062). XC was supported by the USDA NIFA Evans-Allen Grant Numbers NI171445XXXXG004.

## Author Declarations

There is no conflict of interest.

## Authors’ contributions

HLB conceived the research idea. HLB, XC, and DK designed experiments. HS, XC, DK conducted experiments. ZH and HLB analyzed the data. ZH and HLB wrote the manuscript. All authors read and approved the final manuscript.

## Notes

### Competing Interest Statement

The authors have declared no competing interest.

## References

Amdam GV, Page RE. 2010. The developmental genetics and physiology of honeybee societies. Anim Behav 79:973–980.

Amiri E, Strand MK, Rueppell O, Tarpy DR. 2017. Queen quality and the impact of honey bee diseases on queen health: potential for interactions between two major threats to colony health. Insects 8:48.

Baer B, Collins J, Maalaps K, den Boer SP. 2016. Sperm use economy of honeybee (Apis mellifera) queens. Ecology and Evolution 6:2877–2885.

Belsky J, Joshi NK. 2019. Impact of biotic and abiotic stressors on managed and feral bees. Insects 10:233.

Danka RG, Harris JW, Villa JD. 2011. Expression of Varroa sensitive hygiene (VSH) in commercial VSH honey bees (Hymenoptera: Apidae). Journal of economic entomology 104:745–749.

Guichard M, Dainat B, Dietemann V. 2023. Prospects, challenges and perspectives in harnessing natural selection to solve the ‘varroa problem’of honey bees. Evolutionary Applications 16:593–608.

Guzman-Novoa E, Corona M, Alburaki M, Reynaldi FJ, Invernizzi C, Fernández de Landa G, Maggi MD. 2024. Honey bee populations surviving Varroa destructor parasitism in Latin America and their mechanisms of resistance. Frontiers in Ecology and Evolution 12:1434490.

Hinshaw C, Evans KC, Rosa C, López-Uribe MM. 2021. The role of pathogen dynamics and immune gene expression in the survival of feral honey bees. Frontiers in Ecology and Evolution 8:505.

Hunt G, Given JK, Tsuruda JM, Andino GK. 2016. Breeding mite-biting bees to control Varroa. Bee Culture 8:41–47.

Ibrahim A, Reuter GS, Spivak M. 2007. Field trial of honey bee colonies bred for mechanisms of resistance against Varroa destructor. Apidologie 38:67–76.

Kirrane MJ, de Guzman LI, Holloway B, Frake AM, Rinderer TE, Whelan PM. 2015. Phenotypic and genetic analyses of the Varroa sensitive hygienic trait in Russian honey bee (Hymenoptera: Apidae) colonies. PLoS One 10:e0116672.

Kocher SD, Richard F-J, Tarpy DR, Grozinger CM. 2009. Queen reproductive state modulates pheromone production and queen-worker interactions in honeybees. Behavioral Ecology 20:1007–1014.

Kulhanek K, Steinhauer N, Rennich K, Caron DM, Sagili RR, Pettis JS, Ellis JD, Wilson ME, Wilkes JT, Tarpy DR, Rose R, Lee K, Rangel J, vanEngelsdorp D. 2017. A national survey of managed honey bee 2015-2016 annual colony losses in the USA. Journal of Apicultural Research 56:328–340.

Le Conte Y, Ellis M, Ritter W. 2010. Varroa mites and honey bee health: can Varroa explain part of the colony losses? Apidologie 41:353–363.

Lee KV, Goblirsch M, McDermott E, Tarpy DR, Spivak M. 2019. Is the brood pattern within a honey bee colony a reliable indicator of queen quality? Insects 10:12.

López-Uribe MM, Appler RH, Youngsteadt E, Dunn RR, Frank SD, Tarpy DR. 2017. Higher immunocompetence is associated with higher genetic diversity in feral honey bee colonies (Apis mellifera). Conservation Genetics 18:659–666.

McAfee A, Chapman A, Higo H, Underwood R, Milone J, Foster LJ, Guarna MM, Tarpy DR, Pettis JS. 2020. Vulnerability of honey bee queens to heat-induced loss of fertility. Nature Sustain 3:367–376.

Ma C, Zhang Y, Sun J, Imran M, Yang H, Wu J, Zou Y, Li-Byarlay H, Luo S. 2019. Impact of acute oral exposure to thiamethoxam on the homing, flight, learning acquisition and short-term retention of Apis cerana. Pest Management Science 75:2975–2980.

Moore PA, Wilson M E, Skinner JA. 2015. Honey bee queens: evaluating the most important colony member. Bee Health 7(10).

Morfin N, Given K, Evans M, Guzman-Novoa E, Hunt GJ. 2020. Grooming behavior and gene expression of the Indiana “mite-biter” honey bee stock. Apidologie 51:267–275.

Mullin CA, Frazier M, Frazier JL, Ashcraft S, Simonds R, vanEngelsdorp D, Pettis JS. 2010. High levels of miticides and agrochemicals in North American apiaries: implications for honey bee health. PLoS one 5:e9754.

Oddie MAY, Dahle B, Neumann P. 2017. Norwegian honey bees surviving Varroa destructor mite infestations by means of natural selection. Peerj 5.

Phiri BJ, Fèvre D, Hidano A. 2022. Uptrend in global managed honey bee colonies and production based on a six-decade viewpoint, 1961–2017. Sci Rep 12:21298.

Rinderer TE, Harris JW, Hunt GJ, De Guzman LI. 2010. Breeding for resistance to Varroa destructor in North America. Apidologie 41:409–424.

Russo RM, Liendo MC, Landi L, Pietronave H, Merke J, Fain H, Muntaabski I, Palacio MA, Rodríguez GA, Lanzavecchia SB. 2020. Grooming Behavior in Naturally Varroa-Resistant Apis mellifera Colonies From North-Central Argentina. Frontiers in Ecology and Evolution 8:361.

Scaramella N, Burke A, Oddie M, Dahle B, de Miranda J, Mondet F, Rosenkranze P, Neumann P, Locke B. 2023. Host brood traits, independent of adult behaviours, reduce Varroa destructor mite reproduction in resistant honeybee populations. International Journal for Parasitology 53:565–571.

Seeley TD. 2007 Honey bees of the Arnot Forest: a population of feral colonies persisting with Varroa destructor in the northeastern United States. Apidologie 38:19–29.

Siviter H, Fisher A, Baer B, Brown M, Camargo I, Cole J, Le Conte Y, Dorin B, Evans J, Farina W. 2023. Protecting pollinators and our food supply: understanding and managing threats to pollinator health. Insectes Sociaux 1–12.

Smith J, Cleare XL, Given K, Li-Byarlay H. 2021. Morphological Changes in the Mandibles Accompany the Defensive Behavior of Indiana Mite Biting Honey Bees Against Varroa Destructor. Frontiers in Ecology and Evolution 9:243.

Tarpy DR, Caren JR, Delaney DA. 2023. Meta-analysis of genetic diversity and intercolony relatedness among reproductives in commercial honey bee populations. Frontiers in Insect Science 3:5.

Traynor KS, Pettis JS, Tarpy DR, Mullin CA, Frazier JL, Frazier M, Vanengelsdorp D. 2016. In-hive Pesticide Exposome: Assessing risks to migratory honey bees from in-hive pesticide contamination in the Eastern United States. Scientific reports 6:33207.

Tsuruda JM, Harris JW, Bourgeois L, Danka RG, Hunt GJ. 2012. High-Resolution Linkage Analyses to Identify Genes That Influence Varroa Sensitive Hygiene Behavior in Honey Bees. Plos One 7.

Unger P, Guzman-Novoa E. 2010. Maternal effects on the hygienic behavior of Russian× Ontario hybrid honeybees (Apis mellifera L.). Journal of Heredity 101:91–96.

Walsh EM, Sweet S, Knap A, Ing N, Rangel J. 2020. Queen honey bee (Apis mellifera) pheromone and reproductive behavior are affected by pesticide exposure during development. Behavioral Ecology and Sociobiology 74:1–14.

Wang F, Wang Y, Li Y, Zhang S, Shi P, Li-Byarlay H, Luo S. 2022. Pesticide residues in beebread and honey in Apis cerana cerana and their hazards to honey bees and human. Ecotoxicology and environmental safety 238:113574.

Ward K, Cleare X, Li-Byarlay H. 2022. The life span and levels of oxidative stress in foragers between feral and managed honey bee colonies. Journal of Insect Science 22:20.

Warner S, Pokhrel LR, Akula SM, Ubah CS, Richards SL, Jensen H, Kearney GD. 2024. A scoping review on the effects of Varroa mite (Varroa destructor) on global honey bee decline. Sci Total Environ 906: 167492, 10.1016/j.scitotenv.2023.167492.

Wen X, Ma C, Sun M, Wang Y, Xue X, Chen J, Song W, Li-Byarlay H, Luo S. 2021. Pesticide residues in the pollen and nectar of oilseed rape (Brassica napus L.) and their potential risks to honey bees. Science of The Total Environment: 147443.

Winston ML. 1987. THE BIOLOGY OF THE HONEY BEE. Winston, M L the Biology of the Honey Bee Xi+281p Harvard University Press: Cambridge, Massachusetts, USA; London, England, Uk Illus:XI+281P-XI+281P.

Wu-Smart J, Spivak M. 2016. Sub-lethal effects of dietary neonicotinoid insecticide exposure on honey bee queen fecundity and colony development. Scientific reports 6:1–11.

Zheng H, Wang S, Wu Y, Zou S, Dietemann V, Neumann P, Chen Y, Li-Byarlay H, Pirk C, Evans J. 2023. Genomic signatures underlying the oogenesis of the ectoparasitic mite Varroa destructor on its new host Apis mellifera. Journal of advanced research 44:1–11.

